# Suppression of TGF-β/SMAD signaling by an inner nuclear membrane phosphatase complex

**DOI:** 10.1101/2024.09.23.614427

**Authors:** Zhe Ji, Wing-Yan Skyla Siu, Maria Emilia Dueñas, Matthias Trost, Pedro Carvalho

## Abstract

Cytokines of the TGF-β superfamily control essential cell fate decisions via receptor regulated SMAD (R-SMAD) transcription factors. Ligand-induced R-SMAD phosphorylation in the cytosol triggers their activation and nuclear accumulation. We determined how R-SMADs are inactivated by dephosphorylation in the cell nucleus to counteract signaling by TGF-β superfamily ligands. We showed that R-SMAD dephosphorylation is mediated by an inner nuclear membrane associated complex containing the scaffold protein MAN1 and the CTDNEP1-NEP1R1 phosphatase. Structural prediction, domain mapping and mutagenesis revealed that MAN1 binds independently to the CTDNEP1-NEP1R1 phosphatase and R-SMADs to promote their inactivation by dephosphorylation. Disruption of this complex led to nuclear accumulation of R-SMADs and aberrant signaling, even in the absence of TGF-β ligands. These findings establish CTDNEP1-NEP1R1 as the elusive R-SMAD phosphatase and reveal the mechanistic basis for TGF-β signaling inactivation and how this process is disrupted by disease-associated MAN1 mutations.

## Introduction

SMADs constitute a family of transcription factors in animals, essential during embryonic development and adulthood. As key effectors of the transforming growth factor β (TGF-β) superfamily of cytokines, such as TGF-β and Bone Morphogenic Proteins (BMPs), SMADs control a wide range of biological processes like cell differentiation, proliferation, death, adhesion and migration ^1–4^. With such broad and critical roles, it is not surprising that mutations in SMADs proteins have been associated with many diseases including cancer, fibrosis and developmental abnormalities ^5^.

SMAD-dependent gene expression is activated upon the binding of a TGF-β superfamily cytokine to a cognate receptor at the surface of a target cell. Ligand binding stimulates the receptor kinase activity and the subsequent phosphorylation of a receptor-regulated SMAD, or R-SMAD, at conserved serine residues in a C-terminal SXS motif. Once phosphorylated, R-SMADs bind to SMAD4, also known as the common SMAD or Co-SMAD. These SMAD complexes accumulate in the nucleus where they act as transcription factors to regulate gene expression ^2^.

The diversity of transcriptional outputs triggered by TGF-β superfamily cytokines depends largely on the specificity of ligand-receptor interactions and the existence of multiple R-SMADs ^4,6^. For example, while BMP ligands predominantly activate the R-SMADs 1, 5 and 8, whereas TGF-β and activin ligands primarily activate SMAD2 and SMAD3. Thus, despite a common activation mechanism, the diversity of SMAD complexes leads to a wide range of transcriptional outputs.

Activation of R-SMADs by phosphorylation in the cytoplasm is counteracted by dephosphorylation of the C-terminal SXS motif. Based on photobleaching and mathematical modeling experiments, R-SMAD inactivation by dephosphorylation depends on a nuclear localized phosphatase ^7^. Earlier studies suggested that the protein phosphatase Mg2+/Mn2+ dependent 1A (PPM1A) dephosphorylates R-SMADs to terminate signaling by TGF-β superfamily cytokines ^8,9^. However, PPM1A appears to affect R-SMADs indirectly by facilitating their nuclear export ^10^. Consistent with an indirect role, PPM1A knock out (KO) mouse is viable and does not show the developmental defects typical of de-regulated TGF-β/SMAD signaling ^11–13^. Therefore, the identity of the phosphatase and the mechanism of R-SMAD dephosphorylation remains unknown.

CTDNEP1 (C-terminal domain nuclear envelope phosphatase, also known as Dullard) encodes an evolutionarily conserved serine/threonine protein phosphatase localized throughout the endoplasmic reticulum (ER) membrane, including the nuclear envelope ^14,15^. CTDNEP1 works with its regulatory subunit NEP1R1 (nuclear envelope phosphatase 1 regulatory subunit 1), a small ER membrane protein that stabilizes CTDNEP1 ^16,17^.

This CTDNEP1-NEP1R1 phosphatase complex has a well-established and evolutionarily conserved role in dephosphorylating Lipin, a phosphatidic acid hydrolase, thereby contributing to lipid homeostasis ^14,18,19^. Consistent with its role in Lipin regulation, mutations in CTDNEP1-NEP1R1 result in excessive ER membrane proliferation in various cell types ^19,20^. Recently, CTDNEP1-NEP1R1 was also shown to control the stability of SUN2, an inner nuclear membrane (INM) protein important for nuclear organization by transducing cytoskeletal forces into the nucleus ^15,21^.

In mice, deletion of CTDNEP1 results in embryonic lethality due to a defect in formation of primordial germ cells, a process that is critically dependent on signaling by the TGF-β ligand BMP ^22^. The links of CTDNEP1 to TGF-β signaling are supported by additional genetic studies in mice ^23–25^, frogs ^26,27^ and flies ^28–30^. In all cases, loss-of-function mutations in CTDNEP1 result in increased activity of TGF-β/SMAD pathway by unknown mechanisms.

In humans, mutations in CTDNEP1 are frequently observed in aggressive forms of medulloblastoma that display amplification of the C-MYC oncogene ^31–33^. Interestingly, TGF-β signaling is also elevated in these aggressive tumors^33^. The mechanistic basis for the links between CTDNEP1 and TGF-β/SMAD signaling is unknown and cannot be explained by the role of CTDNEP1-NEP1R1 in the dephosphorylation of Lipin, its only substrate described so far.

Here, we showed that CTDNEP1-NEP1R1 is the long-sought phosphatase that dephosphorylates R-SMADs at the C-terminal SXS motif to inactivate signaling by TGF-β ligands. This dephosphorylation occurs in the nucleus and depends on the binding of CTDNEP1-NEP1R1 to the INM protein MAN1, which acts as an R-SMAD-specific adaptor. Disruption of CTDNEP1-NEP1R1-MAN1 complex results in spontaneous nuclear accumulation of R-SMADs and aberrant signaling. These findings provide insight into the mechanism of R-SMAD regulation and explain the molecular basis of diseases and developmental defects caused by unrestrained TGF-β signaling.

## Results

### The CTDNEP1-NEP1R1 phosphatase interacts with the inner nuclear membrane protein MAN1

As a first step in studying the function of CTDNEP1, we searched for its binding partners. CTDNEP1 was fused to a triple FLAG epitope tag (CTDNEP1-FLAG) and expressed in HeLa cells. Like endogenous CTDNEP1 ^15^, CTDNEP1-FLAG localized throughout the ER, including the nuclear envelope (Fig. S1A).

Lysates from cells expressing CTDNEP1-FLAG were prepared in buffer containing 1% DMNG, subjected to immunoprecipitation with anti-FLAG beads and co-precipitated proteins were analyzed by mass spectrometry (Fig. 1A, table S1). Among the most enriched proteins in the CTDNEP1-FLAG precipitates was its regulatory subunit NEP1R1, suggesting that the fusion protein was functional. The CTDNEP1-FLAG precipitates were also highly enriched in MAN1 (also known as LEMD3), a poorly characterized INM resident protein with links to TGFβ/BMP signaling ^34–37^ and mutated in bone disorders characterized by excessive TGFβ/BMP signaling ^38–41^.

**Figure 1.**
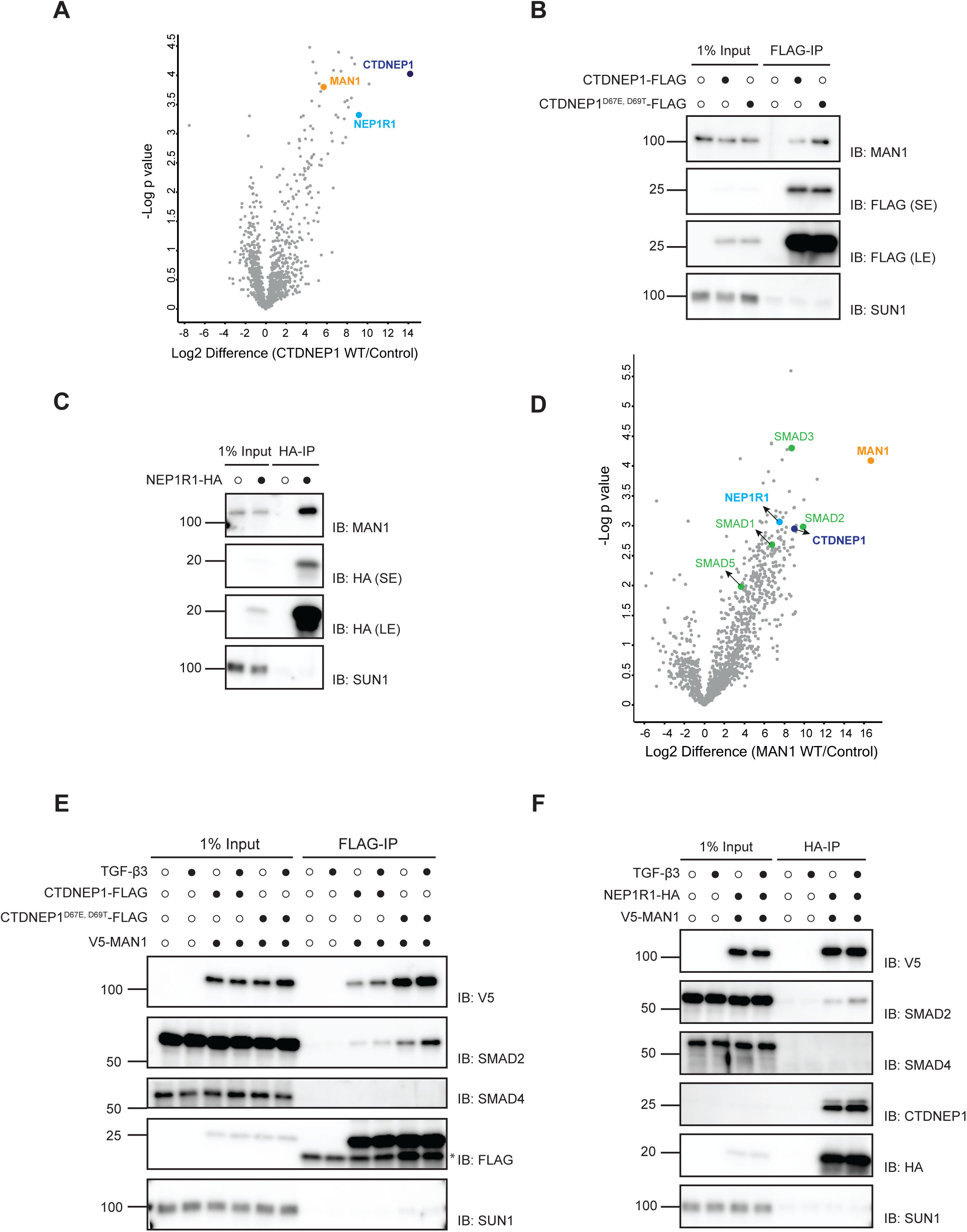
An inner nuclear membrane complex composed of CTDNEP1-NEP1R1-MAN1 interacts with R-SMADs. (A) Proteins co-precipitating with CTDNEP1-FLAG as detected by mass spectrometry. The x-axis shows the log2 fold change of CTDNEP1-FLAG versus untagged control cell line; the y-axis shows the −log10 p-value estimated by the Significance B analysis ^66^. CTDNEP1 partner NEP1R1 is labelled in light blue and MAN1 in orange. (B and C) Immunoprecipitation of FLAG-tagged wild type CTDNEP1 or phosphatase dead CTDNEP1^D67E, D69T^ (B) or NEP1R1-HA (C) from detergent solubilized extracts of HeLa cells. Eluted proteins were analyzed by SDS-PAGE followed by immunoblotting with the indicated antibodies. (D) Proteins co-precipitating with V5-MAN1 as detected by mass spectrometry. The x-axis shows the log2 fold change of V5-MAN1 versus untagged control cell line; the y-axis shows the −log10 p-value estimated by the Significance B analysis ^66^. CTDNEP1 and NEP1R1 are labelled in dark and light blue respectively, and the R-SMADs are labelled in green. (E and F) Immunoprecipitation of CTDNEP1 or CTDNEP1^D67E, D69T^-FLAG (E) or NEP1R1-HA (F) from detergent solubilized extracts of HeLa cells co-expressing V5-MAN1. Immunoprecipitations were performed in absence or upon 1hr treatment with 20ng/ml of TGF-β3. Eluted proteins were analyzed by SDS-PAGE followed by immunoblotting with the indicated antibodies.

The interaction between CTDNEP1-FLAG and MAN1 was confirmed using immunoprecipitation followed by blotting with anti-MAN1 antibodies (Fig. 1B). A mutant that abolishes CTDNEP1 catalytic activity (CTDNEP1^D67E, D69^T) also co-precipitated with MAN1 indicating that the interaction was independent of CTDNEP1 phosphatase activity (Fig. 1B). Similar immunoprecipitations were performed with CTDNEP1’s regulatory subunit, NEP1R1, expressed as a fusion to HA epitope tag (NEP1R1-HA). NEP1R1-HA localized throughout the ER and nuclear envelope, as expected (Fig. S1B). As CTDNEP1, NEP1R1-HA co-precipitated endogenous MAN1 (Fig. 1C).

Conversely, immunoprecipitation of V5-tagged MAN1 (V5-MAN1), which localized to the nuclear rim as was endogenous MAN1 (Fig. S1C), interacted with both CTDNEP1-FLAG (Fig. S1D) and NEP1R1-HA (Fig. S1E). Importantly, interactions between the CTDNEP1-NEP1R1 phosphatase complex and MAN1 appeared specific since SUN1 and other abundant INM proteins were not present in the precipitates, as assayed by immunoblot (Fig. 1B-C, S1D-E) and mass spectrometry (Table S1). Altogether these data indicate that MAN1 is a binding partner of the CTDNEP1-NEP1R1 phosphatase complex, likely at the INM where all three proteins were shown to localize.

### R-SMADs interact with the CTDNEP1-NEP1R1 phosphatase complex

To further explore the role of MAN1 we also analyzed its interactors using V5-MAN1 co-immunoprecipitation followed by mass spectrometry, as described for CTDNEP1. In agreement with our data above, endogenous CTDNEP1 and NEP1R1 were among the most enriched proteins in the V5-MAN1 precipitates (Fig. 1D, Table S1). Other prominent interactors of MAN1 were the R-SMADs 1,2,3 and 5, while no interaction was detected with the co-SMAD SMAD4, as previously observed ^34,35,42^.

These results raised the possibility that the CTDNEP1-NEP1R1 phosphatase complex, MAN1 and R-SMADs were all part of the same protein complex. Consistent with this idea, we observed that CTDNEP1-FLAG (Fig. 1E, S1F) and NEP1R1-HA (Fig. 1F, S1G) coprecipitated both V5-MAN1 and the R-SMADs SMAD1 and 2. In contrast, the Co-SMAD SMAD4 was not detected in the precipitates. Interestingly, CTDNEP1-NEP1R1 interactions with SMAD2 (Fig. 1E, F) and SMAD1 (Fig. S1F, S1G) were observed under basal conditions and were only slightly increased by stimulating SMAD2 and SMAD1 signaling with TGF-β and BMP cytokines, respectively. Together, these data indicate that R-SMADs interact with the CTDNEP1-NEP1R1 phosphatase complex.

### CTDNEP1, NEP1R1 and MAN1 are required for R-SMAD dephosphorylation

R-SMADs are activated by phosphorylation on a conserved SXS motif at their extreme C-termini, resulting in their nuclear accumulation ^1^. How R-SMADs are inactivated by dephosphorylation remains controversial and mysterious ^13,43^. Our finding that R-SMADs form a complex with CTDNEP1-NEP1R1 phosphatase and MAN1 prompted us to test whether these proteins were required for R-SMAD dephosphorylation. A pool of phosphorylated R-SMADs was generated by stimulating cells with TGF-β or BMP, and the kinetics of R-SMAD dephosphorylation and R-SMAD localization were analyzed after washing out the activating ligands. Inhibitor of TGF-β or BMP cell surface receptors were also added to prevent R-SMAD re-phosphorylation (Fig. 2A), as previously described ^8^. In parental HeLa, phospho-SMAD2 had a half-life of less than 60 minutes. Similarly, deletion of the phosphatase PPM1A, previously implicated in SMAD dephosphorylation ^8^, did not affect the kinetics of SMAD2 dephosphorylation (Fig. 2B, S2A). In contrast, loss of MAN1, CTDNEP1 or NEP1R1 strongly inhibited SMAD2 dephosphorylation without affecting the overall levels of SMAD2 protein (Fig. 2B, S2A). Depletion of MAN1, CTDNEP1 or NEP1R1 also resulted in delayed SMAD2 dephosphorylation in U2OS cells (Fig. S2B). Consistent with delayed R-SMAD dephosphorylation, MAN1, CTDNEP1 and NEP1R1 KO cells showed persistent SMAD2 nuclear accumulation 7 hours upon TGF-β inhibition, while in control cells SMAD2 nuclear accumulation largely dissipated within 1 hour after TGF-β inhibition (Fig. S2C, D). Moreover, MAN1, CTDNEP1 and NEP1R1 KO cells also displayed slowed kinetics of SMAD1/5/8 dephosphorylation after activation with BMP (Fig. 2C). Altogether these data indicate that upon stimulation with TGF-β superfamily ligands, R-SMAD dephosphorylation requires MAN1, CTDNEP1 and NEP1R1.

**Figure 2.**
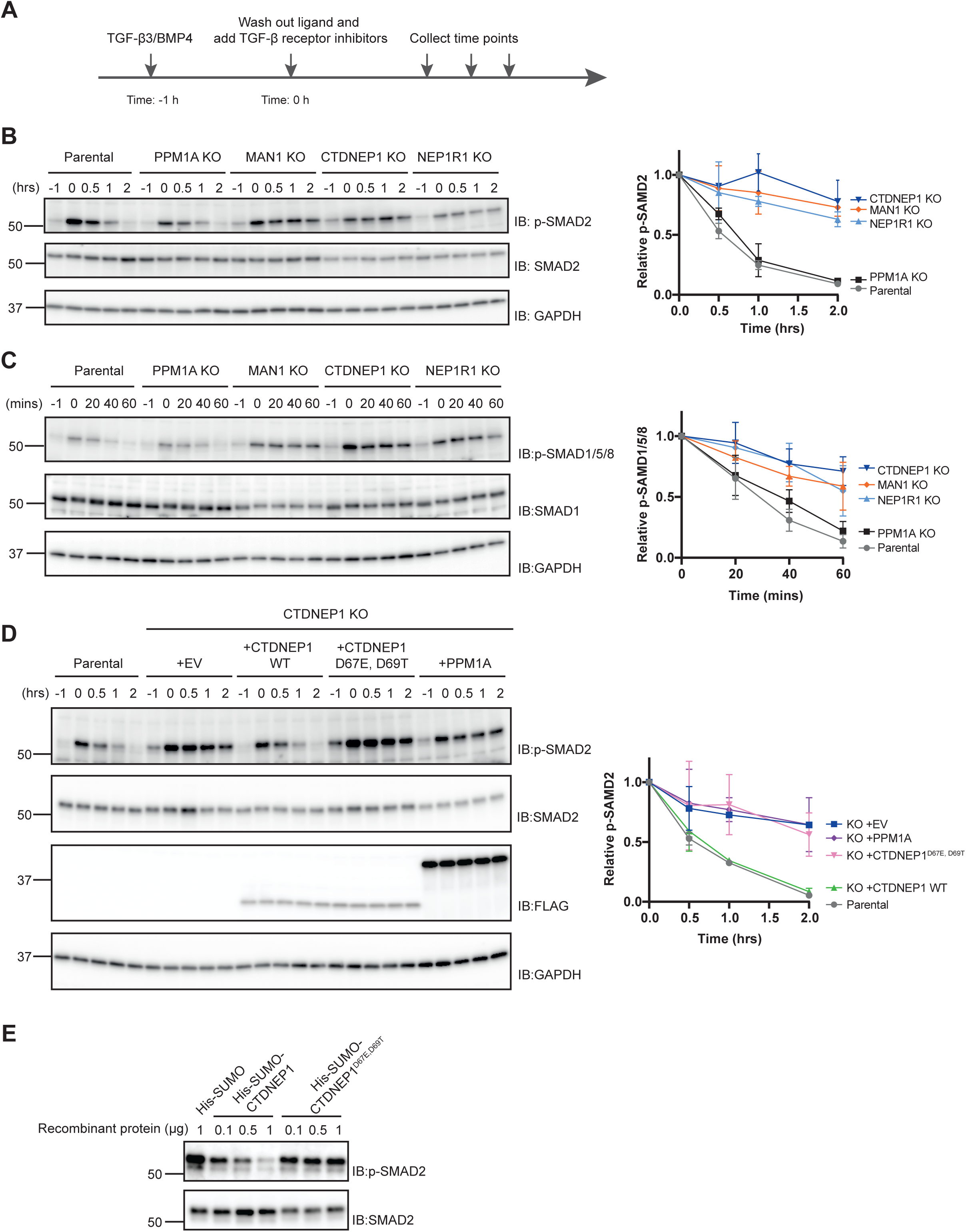
R-SMAD dephosphorylation requires MAN1 and the CTDNEP1-NEP1R1 phosphatase. (A) Scheme of the experimental outline to monitor the kinetics of R-SMAD dephosphorylation. (B) Time course analysis of SMAD2 dephosphorylation upon TGF-β3 stimulation in parental, PPM1A, MAN1, CTDNEP1 and NEP1R1 KO HeLa cells. Cell extracts were analyzed by SDS–PAGE and immunoblotting with the indicated antibodies. The graph (right) shows the average of three experiments; error bars represent the standard deviation. (C) Time course analysis of SMAD1/5/8 dephosphorylation upon BMP4 stimulation in parental, PPM1A, MAN1, CTDNEP1 and NEP1R1 KO HeLa cells. Cell extracts were analyzed by SDS–PAGE and immunoblotting with the indicated antibodies. The graph (right) shows the average of three experiments; error bars represent the standard deviation. (D) Time course analysis of SMAD2 dephosphorylation upon TGF-β3 stimulation in parental and CTDNEP1 KO HeLa cells expressing the indicated proteins. Cell extracts were analyzed by SDS–PAGE and immunoblotting with the indicated antibodies. The graph (right) shows the average of two experiments; error bars represent the standard deviation. (E) Analysis of SMAD2 dephosphorylation by recombinant wild type CTDNEP1 and phosphatase dead CTDNEP1^D67E, D69T^ expressed as fusion proteins to a His-SUMO tag. Purified His-SUMO was also used as a negative control. Note that soluble versions of CTDNEP1 and of CTDNEP1^D67E,D69T^ were generated by deleting the N terminal amphipathic helix of CTDNEP1 corresponding to amino acids 1-45.

Using the assay described above, we tested whether CTDNEP1 phosphatase activity was necessary for R-SMAD dephosphorylation. We observed that re-expression of wild type CTDNEP1 in CTDNEP1 KO cells restored normal kinetics of SMAD2 dephosphorylation. In contrast, expression of catalytically inactive CTDNEP1^D67E, D69T^ failed to promote SMAD2 dephosphorylation (Fig. 2D). Importantly, CTDNEP1^D67E, D69T^ is expressed to normal levels and appears to interact normally with its partners, including SMAD2 (Fig. 1E). Thus, CTDNEP1 phosphatase activity is essential for R-SMAD dephosphorylation. Moreover, overexpression of the phosphatase PPM1A was unable to compensate for the absence of CTDNEP1 indicating that its function in R-SMAD dephosphorylation is specific (Fig. 2D).

Next, we asked if CTDNEP1 was able to directly dephosphorylate R-SMADs. Purified phospho-SMAD2 immobilized on beads was incubated with recombinant CTDNEP1 or CTDNEP1^D67E, D69T^ expressed as fusion to a His-SUMO tag (Fig. S3A). While SMAD2 efficiently dephosphorylated by CTDNEP1 in a concentration dependent manner, CTDNEP1^D67E, D69T^ was unable to do so even when present at high concentrations (Fig. 2E). Collectively, these experiments indicate that R-SMAD dephosphorylation requires INM protein MAN1 and the CTDNEP1-NEP1R1 phosphatase.

### MAN1 is a R-SMAD adaptor for the CTDNEP1-NEP1R1 phosphatase

Our data are consistent with a model in which the CTDNEP1-NEP1R1-MAN1 complex functions as the elusive R-SMAD phosphatase. To explore this possibility, we investigated how interactions among CTDNEP1, NEP1R1 and MAN1 contributed to R-SMAD dephosphorylation.

Given that MAN1 was shown to bind R-SMADs directly ^44^, a simple possibility was that MAN1 bridged the interaction between the R-SMADs and the CTDNEP1-NEP1R1 phosphatase complex. Indeed, in MAN1 KO cells, both CTDNEP1-FLAG (Fig. 3A) and NEP1R1-HA (Fig. 3B) failed to interact with the SMAD1 and SMAD2, even upon TGF-β stimulation that leads to their nuclear accumulation and activation of SMAD signaling. Importantly, loss of MAN1 did not affect the localization of CTDNEP1 (Fig. 3C) or its interaction with NEP1R1 (Fig. 3B). To test whether loss of MAN1 had a general effect on CTDNEP1-NEP1R1 activity we analyzed the levels of Lipin1 and SUN2, two proteins with reduced steady state levels in the absence of CTDNEP1-NEP1R1 phosphatase ^19,21,45^. In contrast with CTDNEP1 and NEP1R1 KO cells, MAN1 KO HeLa cells display normal levels of both Lipin 1 and SUN2 indicating that MAN1 is not a general regulator of CTDNEP1-NEP1R1 activities and regulates selectively R-SMADs dephosphorylation (Fig. S3B). Together these data suggest that MAN1 acts as INM scaffold facilitating specifically the interaction between R-SMADs and the CTDNEP1-NEP1R1 phosphatase complex.

**Figure 3.**
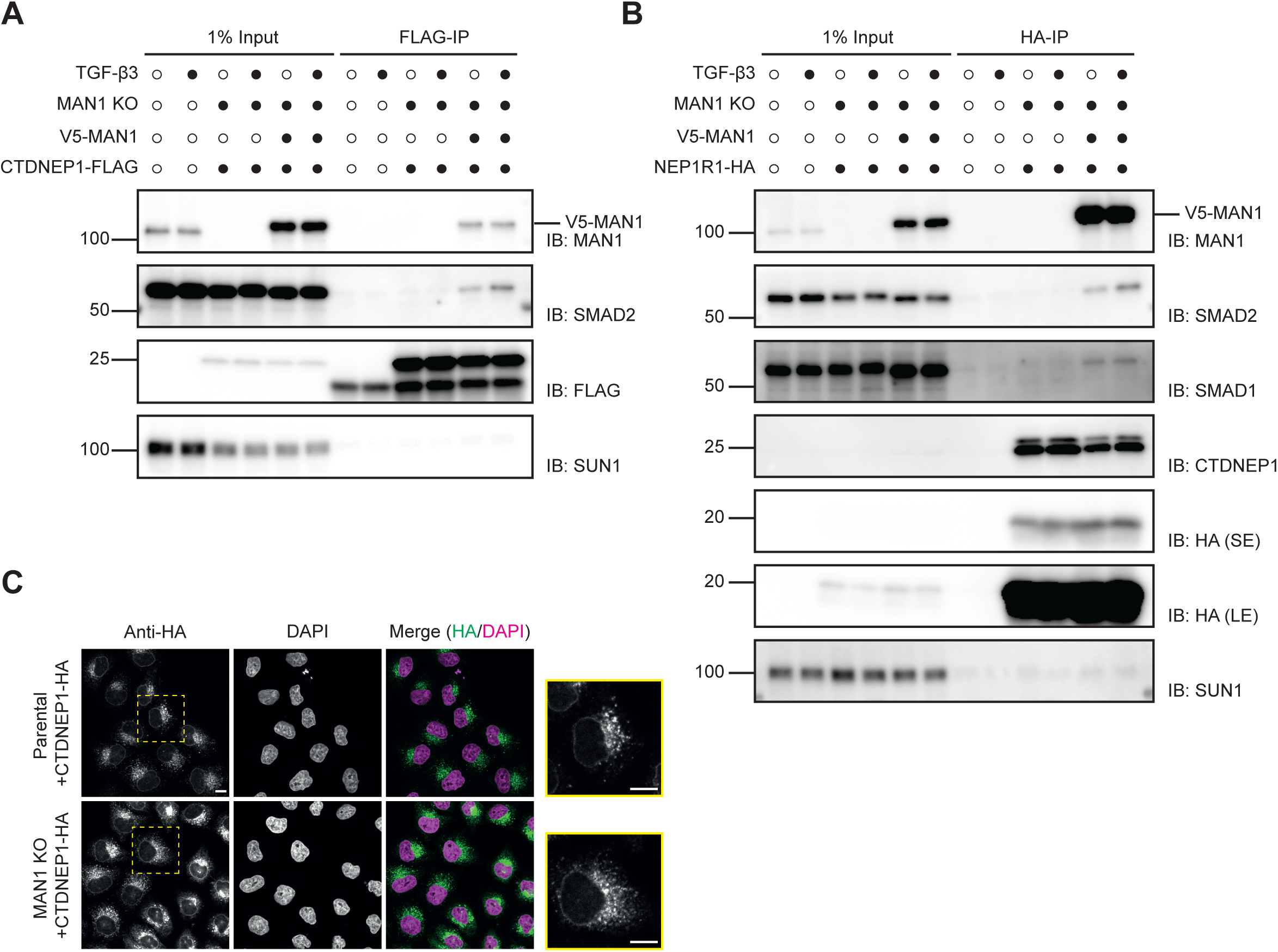
MAN1 is required for the interaction between CTDNEP1- NEP1R1 phosphatase and R-SMADs. (A) Immunoprecipitation of FLAG-tagged CTDNEP1 from detergent solubilized extracts of MAN1 KO HeLa cells transduced with an empty vector or a construct encoding V5-MAN1. Immunoprecipitations were performed in the absence or upon a 1hr treatment with 20ng/mL of TGF-β3, as indicated. Eluted proteins were analyzed by SDS-PAGE followed by immunoblotting with the indicated antibodies. (B) Immunoprecipitation of HA-tagged NEP1R1 from detergent solubilized extracts of MAN1 KO HeLa cells transduced with an empty vector or a construct encoding V5-MAN1. Immunoprecipitations were performed in the absence or upon a 1hr treatment with 20ng/ml of TGF-β3, as indicated. Eluted proteins were analyzed by SDS-PAGE followed by immunoblotting with the indicated antibodies. (C) Localization of CTDNEP1-HA in parental and MAN1 KO HeLa cells analyzed by immunofluorescence. CTDNEP1-HA was detected with anti-HA antibodies and DNA was labelled with 4’,6- diamidino- 2- phenylindole (DAPI). Dotted yellow square indicates the zoomed in cell shown on the far right. Scale bar:10μM.

A prediction of this model was that MAN1 would interact independently with R-SMADs and the CTDNEP1-NEP1R1 phosphatase complex. We explored this possibility by the interactions between MAN1 and the CTDNEP1-NEP1R1 phosphatase complex using immunoprecipitation. Interestingly, regulatory and catalytic subunits showed distinct dependencies in their binding to MAN1. While the interaction between CTDNEP1 with MAN1 required NEP1R1 (Fig. 4A), the binding of NEP1R1 to MAN1 was only slightly reduced in CTDNEP1 KO cells (Fig. 4B) suggesting a central role of NEP1R1 in the recruitment of the CTDNEP1-NEP1R1 phosphatase complex to MAN1. In agreement with these findings, structural analysis using Alphafold multimer predicted an interaction between NEP1R1 and MAN1 through their membrane regions (Fig. S4A, B) ^46,47^. To gain further insight into the NEP1R1-MAN1 binding mechanism, we tested these predictions by generating chimeric proteins in which either one or both of MAN1 transmembrane segments were replaced by the ones of its paralogue LEMD2 (Fig. 4C) ^48^. Despite a similar domain organization and inner nuclear membrane localization, LEMD2 functions primarily in maintaining nuclear envelope integrity, by facilitating its reassembly during mitosis and repair upon damage ^49–51^. Remarkably, both MAN1^LEMD2TM1^ and MAN1^LEMD2TM1+2^ failed to interact with NEP1R1 (Fig. 4D).

**Figure 4.**
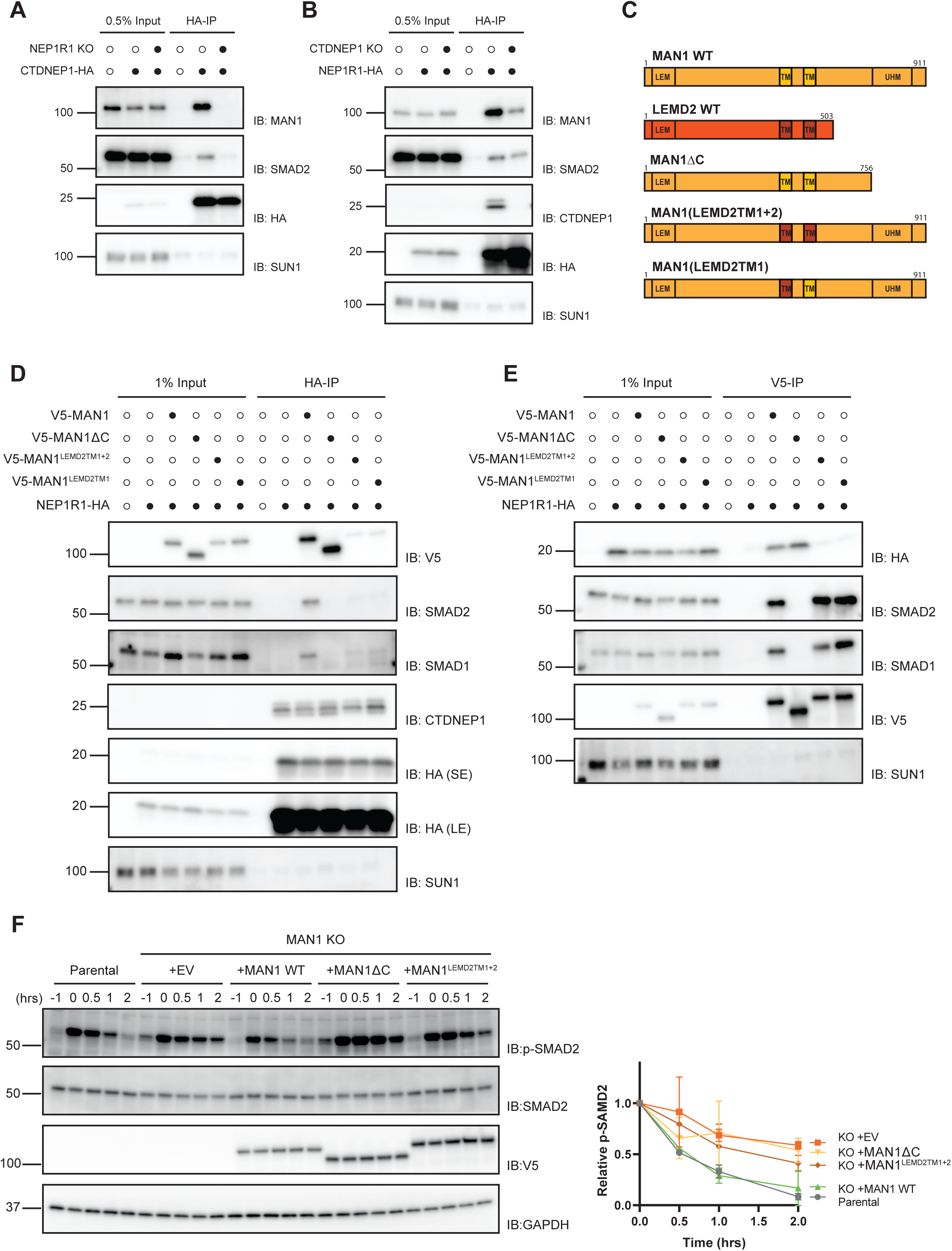
MAN1 binds CTDNEP1-NEP1R1 phosphatase and R-SMADs independently. (A) Immunoprecipitation of HA-tagged CTDNEP1 from detergent solubilized extracts in parental or NEP1R1 KO HeLa cells. Eluted proteins were analyzed by SDS-PAGE followed by immunoblotting with the indicated antibodies. (B) Immunoprecipitation of HA-tagged NEP1R1 from detergent solubilized extracts in parental or CTDNEP1 KO HeLa cells. Eluted proteins were analyzed by SDS-PAGE followed by immunoblotting with the indicated antibodies. (C) Schematic representation of MAN1 and related proteins. (D) Immunoprecipitation of HA-tagged NEP1R1 from detergent solubilized extracts of MAN1 KO HeLa cells transduced with an empty vector or the indicated V5-tagged MAN1 derivatives. Eluted proteins were analyzed by SDS- PAGE followed by immunoblotting with the indicated antibodies. (E) Immunoprecipitation of the indicated V5-tagged MAN1 derivatives from detergent solubilized extracts of MAN1 KO HeLa cells co-expressing HA- tagged NEP1R1. Eluted proteins were analyzed by SDS-PAGE followed by immunoblotting with the indicated antibodies. (F) Time course analysis of SMAD2 dephosphorylation upon TGF-β3 stimulation in parental and MAN1 KO HeLa cells expressing the indicated proteins. Cell extracts were analyzed by SDS–PAGE and immunoblotting with the indicated antibodies. The graph (right) shows the average of two experiments; error bars represent the standard deviation.

Like MAN1, the MAN1^LEMD2TM^ mutants localized to the nuclear envelope (Fig. S4C) and interacted normally with R-SMADs (Fig. 4E). Therefore, MAN1 interaction with the CTDNEP1-NEP1R1 phosphatase complex depends on the binding between NEP1R1 and MAN1 transmembrane regions.

Next, we examined the contribution of MAN1 C-terminal region for the interactions with the phosphatase complex and R-SMADs. MAN1 C-terminus encompasses the UHM domain previously shown to bind to R-SMADs ^34,35,44^. We observed that MAN1ΔC, a MAN1 truncation lacking its last 155 amino acids including the UHM domain, still localized to the INM (Fig. S4C) but failed to interact with R-SMADs (Fig. 4E), as expected. On the other hand, MAN1ΔC efficiently interacted with the CTDNEP1-NEP1R1 phosphatase complex (Fig. 4D, E). Thus, MAN1 functions as a scaffold using distinct domains to interact with R-SMADs and CTDNEP1-NEP1R1 phosphatase complex.

Finally, we assessed the importance of these MAN1 domains for SMAD2 dephosphorylation. The SMAD2 dephosphorylation defect in MAN1 KO cells was reversed by expression of wild-type MAN1 but not by expression of MAN1^LEMD2TM1+2^ or MAN1ΔC, impaired in CTDNEP1-NEP1R1 and R-SMAD binding, respectively (Fig. 4F). Since these MAN1 derivatives display similar levels and localization (Fig. S4C, D), we concluded that MAN1-dependent R-SMAD dephosphorylation requires its binding both to the CTDNEP1-NEP1R1 and R-SMADs. Together, these results indicate that MAN1 functions as an R-SMAD-specific adaptor for the CTDNEP1-NEP1R1 phosphatase at the INM.

### CTDNEP1, NEP1R1 and MAN1 suppress inappropriate SMAD signaling

Activation of R-SMADs with TGF-β family cytokines leads to their phosphorylation and nuclear accumulation ^1,52^. Intriguingly, depletion of CTDNEP1, NEP1R1 or MAN1 triggered nuclear accumulation of SMAD2 even in the absence of exogenous TGF-β ligands (Fig. 5A, B). This effect appeared specific as SMAD2 distribution was unaffected by depletion of the phosphatase PPM1A (Fig. 5A, B). The normal SMAD2 distribution was restored upon re-expression of the wild type in the corresponding KO cells (Fig. S5A, B). On the other hand, in CTDNEP1 KO cells, SMAD2 nuclear accumulation was not reversed by expression of phosphatase dead CTDNEP1 or by active PPM1A, highlighting the importance of CTDNEP1 catalytic activity in regulating SMAD2 distribution (Fig. S5A, B). Given that R-SMADs nuclear localization requires their prior phosphorylation at the C-terminal SXS motif by receptor activated kinases, we tested if in the absence of exogenous TGF-β ligands, CTDNEP1, NEP1R1 or MAN1 KO cells showed increased SMAD2 phosphorylation. Indeed, under these basal conditions, the KO cells showed increased levels of phosphorylated SMAD2 which was undetectable in control cells (Fig. 5C). Importantly, SMAD2 phosphorylation in CTDNEP1 KO cells depended on TGF-β receptor kinase activity as it was reversed by the specific kinase inhibitor SB431542 (Fig. 5D).

**Figure 5.**
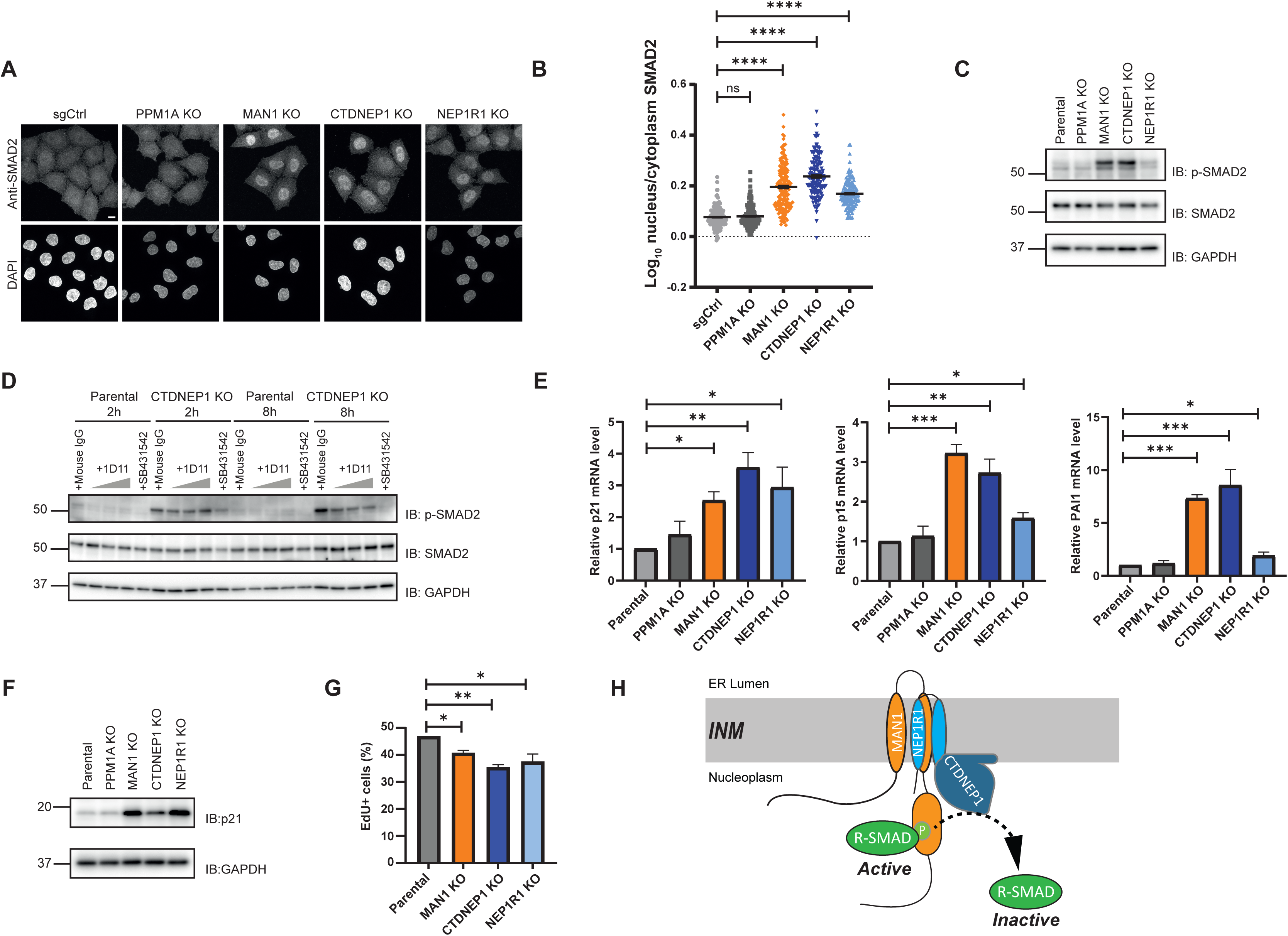
The CTDNEP1-NEP1R1 phosphatase and MAN1 suppress inappropriate SMAD signaling. (A) localization of endogenous SMAD2 in HeLa parental cells or lacking the indicated genes analyzed by immunofluorescence. DNA was labelled with 4’,6- diamidino- 2- phenylindole (DAPI). Scale bar:10μM. (B) Quantification of nuclear accumulation of SMAD2 from imaging experiments as shown in (A) from three replicates. Error bars represent the standard error mean of the three replicates. n = 100, **** p<0.0001. (C) levels of endogenous p-SMAD2 and SMAD2 in HeLa parental cells or lacking the indicated genes. Cell lysates were analyzed by SDS-PAGE followed by immunoblotting with the indicated antibodies. GAPDH was used as loading control. (D) levels of endogenous p-SMAD2 and SMAD2 in HeLa parental cells or CTDENP1 KO cells with indicated treatments. Mouse IgG antibody was used at 300 µg/mL, 1D11 antibody was used at 30, 150 and 300 µg/mL respectively. Cell lysates were analysed by SDS-PAGE followed immunoblotting with the indicated antibodies. GAPDH was used as loading control. (E) levels of p21, p15 and PAI1 transcripts in HeLa parental cells or lacking the indicated genes analyzed by RT-qPCR. n = 2, * p<0.05, ** p<0.01, *** p<0.001. (F) levels of endogenous p21 in HeLa parental cells or lacking the indicated genes. Cell lysates were analyzed by SDS-PAGE followed by immunoblotting with the indicated antibodies. GAPDH was used as loading control. (G) Quantification of S phase cells assessed based on EdU incorporation and DAPI staining. At least 30000 cells were analyzed for each condition. n = 2, * p<0.05, ** p<0.01 (H) The CTDNEP1-NEP1R1-MAN1 complex dephosphorylates and inactivates R-SMADs at the INM (see text for details).

Increased levels of phosphorylated SMAD2 in the KO cells could result from autocrine signaling due to increased production of TGB ligands, as previously described ^53,54^. Consistently, MAN1, CTDNEP1 and NEP1R1KO cells expressed higher levels of TGF-β1 and 2 ligands, as detected by RT-qPCR (Fig S5D). The addition of a neutralizing antibody 1D11, which specifically binds to and inhibits TGF-β ligands ^55^, to the cell media reduced the levels of phosphorylated SMAD2 in CTDNEP1KO cells further suggesting they have aberrant autocrine TGF-β signalling (Fig 5D, S5C). However, the neutralizing antibody, even when added in large excess, could not efficiently suppress the SMAD2 phosphorylation in CTDNEP1KO cells, as observed with the TGF-β receptor kinase inhibitors SB431542 (Fig 5D). These data suggested that TGF-β receptor has some basal kinase activity, normally counteracted by the phosphatase activity of CTDNEP1.

Next, we asked if the accumulation of phosphorylated SMAD2 in the KO cells was sufficient to trigger expression of several SMAD target genes (Fig. 5E), including PAI1 and the cell cycle inhibitors p15, p21 (Fig. 5E, F). Importantly, the accumulation of cell cycle inhibitors was functionally relevant as the KO cells displayed growth inhibition in the absence of TGF-β ligands (Figure 5G).

Therefore, besides the essential role in R-SMAD inactivation upon TGF-β ligands stimuli, the CTDNEP1-NEP1R1-MAN1 complex also has a critical and constitutive function in suppressing aberrant SMAD signaling, due to both autocrine and basal activation of the TGF-β receptor.

## Discussion

Accurate signaling via TGF-β superfamily cytokines depends on tight regulation of SMAD activity. While the mechanisms of SMAD activation by phosphorylation have been extensively characterized, much less is known about their inactivation. Here, we determined the mechanism of R-SMAD inactivation by dephosphorylation and identified CTDNEP1-NEP1R1 as the long-sought R-SMAD phosphatase.

We showed that the dephosphorylation of R-SMAD requires their interaction with an INM complex composed of MAN1 and the phosphatase CTDNEP1-NEP1R1. In this complex, MAN1 functions as a scaffold, bridging the interaction between the CTDNEP1-NEP1R1 phosphatase and phosphorylated R-SMADs thereby promoting their dephosphorylation (Fig. 5G). The interaction with R-SMADs requires MAN1 C-terminal nucleoplasmic domain, consistent with earlier studies ^34,42,44^. MAN1 interaction with the phosphatase complex occurs via the regulatory subunit NEP1R1 and involves the transmembrane regions of both proteins. Thus, besides a general role in stabilizing CTDNEP1 ^16^, NEP1R1 is critical in SMAD regulation by promoting the interaction of the phosphatase complex with MAN1.

The regulation of other CTDNEP1-NEP1R1 substrates, such as Lipin ^19^ and the recently identified SUN2 ^21^, does not appear to require MAN1. Instead, MAN1 functions as a specific and essential adaptor for R-SMADs dephosphorylation. MAN1 exclusive localization to the INM ensures that R-SMAD dephosphorylation occurs only in the nucleus. These findings provide the mechanistic basis to explain previous results from mathematical modelling and time-course experiments proposing that precise regulation of TGF-β signaling required R-SMADs to be dephosphorylated in the nucleus but not in the cytosol ^7,56^.

In active SMAD complexes, the phosphorylated SXS motif of one R-SMAD interacts with a basic pocket of a partner SMAD protein, either another R-SMAD or the co-SMAD SMAD4 ^57,58^. Thus, exposure of the SXS motif for dephosphorylation by the CTDNEP1-NEP1R1-MAN1 complex likely occurs in a coordinated fashion with SMAD complex disassembly. Monoubiquitination of SMADs, in particular SMAD3 and SMAD4, was shown to stimulate the disassembly of SMAD complexes ^59–61^. In the case of SMAD3, ubiquitination is mediated by the ubiquitin ligase SMURF2 and occurs at a lysine residue conserved in other R-SMADs suggesting that this may be part of general mechanism to disassemble SMAD complexes ^61,62^. Monoubiquitination of SMAD4 involves a different ubiquitin ligase, TRIM33 ^59,60^. Curiously, the binding of TRIM33 to specific chromatin domains stimulates its ubiquitination activity towards SMAD4 providing an additional level of regulation of SMAD complex disassembly ^63^. In the future it will be interesting to test whether SMAD complex disassembly by monoubiquitination and CTDNEP1-NEP1R1-MAN1-dependent R-SMAD dephosphorylation are coordinated and if this potential coordination contributes for the temporal and spatial inactivation of SMAD signaling.

Binding of a TGF-β family cytokine to a cognate receptor at the surface of a target cell is known to initiate SMAD signaling ^1,3^. We found that disruption of the CTDNEP1-NEP1R1-MAN1 complex not only delays SMAD inactivation upon ligand-induced stimulation but also triggers aberrant SMAD signaling in the absence of exogenous ligands. This observation suggests that, like other kinases, the intrinsic basal activity of TGF-β cytokine receptors is sufficient to phosphorylate and activate SMADs. Our data indicates that this activity is normally counteracted by the CTDNEP1-NEP1R1-MAN1 complex that constitutively dephosphorylate R-SMADs. In agreement with this model, we observe that CTDNEP1-NEP1R1-MAN1 complex is present in cells and interacts with R-SMADs irrespective of TGF-β ligand stimulation. Moreover, the levels and composition of the CTDNEP1-NEP1R1-MAN1 complex appeared unchanged upon ligand-induced stimulation of TGF-β signaling further supporting that it functions constitutively.

The mechanism of R-SMAD inactivation determined here also provides the molecular framework to understand the plethora of phenotypes caused by CTDNEP1 and MAN1 mutations in a variety of model organisms, such as flies, frogs and mice ^23–29,34,36,37,42^. Whole body deletion of CTDNEP1 is embryonic lethal in mice ^22^, while its conditional ablation leads to strong SMAD signalling deregulation in various tissues including kidney ^23^, bone ^64^, heart ^25^ and ovaries ^65^. Moreover, MAN1 mutations result in a variety of disorders characterized by increased bone density, such as osteopoikilosis, Buschke-Ollendorff syndrome and melorheostosis ^38–41^. While the common denominator among these phenotypes is an increase in TGF-β/BMP signaling, the roles of CTDNEP1 and MAN1 remained enigmatic. This is now illuminated by our study. We also provide insight into the function of NEP1R1, for which there was less information aside from promoting CTDNEP1 stability ^16,17^.

Aggressive medulloblastomas frequently display CTDNEP1 mutations ^31,32^. Loss of CTDNEP1 appears to potentiate amplification of the MYC oncogene ^33^. It was suggested that dephosphorylation of MYC Serine 62 by CTDNEP1 curbs its oncogenic activity. Interestingly, loss of CTDNEP1 also correlated with increased TGF-β signaling in these tumors. In the future, it will be interesting to explore if these two activities of CTDNEP1 are linked and how they contribute to medulloblastoma progression.

The role of CTDNEP1-NEP1R1 phosphatase in the regulation of Lipin has received great attention. Our study shows that this phosphatase has more pervasive functions in cell regulation. Other recent studies also suggest that CTDNEP1-NEP1R1 display additional substrates ^15,20,21^. In the future, it will be important to identify the complete substrate set of this phosphatase and understand their regulation.

## Acknowledgements

We thank C. Hill for discussions and U. Gruneberg, C. Hill and R. Klemm for critical reading of the manuscript. PC was supported by an investigator award from the Wellcome Trust (223153/Z/21/Z) and an ERC consolidator grant (GA 817708). This research was co-funded by grant awards to MT (Wellcome Trust Multi-User Equipment grant (212947/Z/18/Z) and Investigator Award (215542/Z/19/Z)).

## Author Contributions

Z.J. and W.S.S. performed all the experiments. Z.J. and W.S.S. analyzed the data with the help of P.C.. M.E.D. and M.T. generated and analyzed the mass spectrometry data. P.C. conceived and supervised the project and wrote the manuscript with input from all the authors.

## Declaration of Interests

The authors declare no competing interests.

## Legends to the supplementary figures

**Figure S1. Localization and immunoprecipitation analysis of CTDNEP1, NEP1R1 and MAN1**

Localization of CTDNEP1-FLAG, NEP1R1-HA (B) and V5-MAN1 (C) in HeLa cells analyzed by immunofluorescence. CTDNEP1, NEP1R1 and MAN1 were detected with anti-FLAG, -HA and –V5 antibodies, respectively and DNA was labelled with 4’,6-diamidino-2-phenylindole (DAPI). Note that anti-FLAG antibody non-specifically labels the cell periphery. Scale bar:10μM.

(D and E) Immunoprecipitation of V5-MAN from detergent solubilized extracts of HeLa cells co-expressing either CTDNEP1 or CTDNEP1^D67E, D69T –^ FLAG (D) or NEP1R1-HA (E). Eluted proteins were analyzed by SDS-PAGE followed by immunoblotting with the indicated antibodies.

(F and G) Immunoprecipitation of CTDNEP1 or CTDNEP1^D67E, D69T^-FLAG (F) or NEP1R1-HA (G) from detergent solubilized extracts of HeLa cells co-expressing V5-MAN1. Immunoprecipitations were performed in absence or upon 1hr treatment with 20ng/ml of BMP4. Eluted proteins were analyzed by SDS-PAGE followed by immunoblotting with the indicated antibodies.

**Figure S2. R-SMAD dephosphorylation requires MAN1 and the CTDNEP1-NEP1R1 phosphatase in U2OS cells**

(A) Validation of PPM1A, MAN1, CTDNEP1 and NEP1R1 knock out HeLa cells. Cell extracts were analyzed by SDS-PAGE followed by immunoblotting with the indicated antibodies.

(B) Time course analysis of SMAD2 dephosphorylation upon TGF-β3 stimulation in parental, PPM1A, MAN1, CTDNEP1 and NEP1R1 KO U2OS cells.

Cell lysates were subjected to SDS-PAGE separation and immunoblotting was performed with the indicated antibodies. The graph (right) shows the average of three experiments; error bars represent standard deviation.

(C) Immunofluorescence of time course analysis of endogenous SMAD2 localization upon TGF-β3 stimulation in parental, MAN1 KO, CTDNEP1 KO and NEP1R1 KO HeLa cells.

(D) Quantification of nuclear accumulation of SMAD2 from imaging experiments as shown in (C) from two independent biological replicates. Error bars represent the standard error mean of the two replicates (n > 60).

**Figure S3. R-SMAD dephosphorylation requires CTDNEP1-NEP1R1 phosphatase**

(A) Purified His-SUMO tagged wild type or phosphatase dead CTDNEP1, or His-SUMO alone. The purity of the recombinantly expressed proteins was analyzed by SDS-PAGE followed by staining with Coomassie blue. Note that soluble versions of CTDNEP1 and of CTDNEP1^D67E,D69T^ were generated by deleting the N terminal amphipathic helix of CTDNEP1 corresponding to amino acids 1-45.

(B) Steady state levels of Lipin and SUN2 are specifically affected by the loss of CTDNEP1 and NEP1R1 while loss of MAN1 has no effect. Extracts of HeLa cells with the indicated genotype were analyzed by SDS-PAGE followed by immunoblotting with anti-Lipin1 and anti-SUN2 antibodies. GAPDH was used as a loading control and detected with an anti-GAPDH antibody.

**Figure S4. Different MAN1 domains interact with NEP1R1 and R-SMADs**

(A) AlphaFold multimer structural model of NEP1R1 (light blue) and MAN1 (orange).

(B) Alphafold predicted alignment error (PAE) plot of the model shown in (A). The predicted interaction between MAN1 and NEP1R1 membrane regions is indicated by the dotted yellow box.

(C) Localization of endogenous SMAD2 in HeLa parental and MAN1 KO cells expressing the indicated V5-tagged MAN1 derivatives analyzed immunofluorescence. SMAD2 and MAN1 derivatives were detected with anti-SMAD2 and anti-V5 antibodies, respectively. DNA was labelled with 4’,6-diamidino-2-phenylindole (DAPI). Scale bar:10μM

(D) Quantification of nuclear accumulation of SMAD2 from imaging experiments as shown in (C) from three replicates. Error bars represent the standard error mean of the three replicates. n = 100, * p<0.05, **** p<0.0001

**Figure S5. CTDNEP1, NEP1R1 suppress aberrant SMAD signaling**

(A) Localization of endogenous SMAD2 in HeLa parental, CTDNEP1 KO or NEP1R1 KO cells expressing the indicated HA-tagged proteins analyzed immunofluorescence. SMAD2 was detected with an anti-SMAD2 antibody and CTDNEP1 derivatives and NEP1R1 were detected with an anti-HA antibody. DNA was labelled with 4’,6-diamidino-2-phenylindole (DAPI). Scale bar:10μM

(B) Quantification of nuclear accumulation of SMAD2 from imaging experiments as shown in (A) from three replicates. Error bars represent the standard error mean of the three replicates. n = 100, **** p<0.0001

(C) Levels of endogenous p-SMAD2 and SMAD2 in HeLa parental cells with indicated treatments. Mouse IgG antibody was used at 300 µg/mL, 1D11 antibody was used at 30, 150 and 300 µg/mL respectively. Cell lysates were analysed by SDS-PAGE followed immunoblotting with the indicated antibodies. GAPDH was used as loading control.

(D) Levels of TGF-β1, 2 and 3 transcripts in HeLa parental cells or lacking the indicated genes analyzed by RT-qPCR. n = 2, * p<0.05, ** p<0.01.

## Material and Methods

### Cells

HeLa cells were obtained from the ATCC. U2OS cells were obtained from the ECACC. The Lenti-X 293T cell line for production of lentivirus was obtained from TakaraBio. Flp-In T-REx HEK293 cells were obtained from Invitrogen (Thermo Fischer Scientific). All cells were grown at 37°C 5% CO2 in DMEM medium (Merck Life Science UK Limited #D6429) supplemented with L-Glutamine (2 mM; Gibco #25030024), Penicillin-Streptomycin (10 Units/mL; Gibco #15140122) and 10% FCS (Merck Life Science UK Limited #F9665).

For expressing recombinant CTDNEP1 proteins for the in vitro phosphatase assay, BL21-CodonPlus (DE3)-RIPL Competent cells (Agilent Technologies #230280) were cultured at 37°C in LB media containing Kanamycin and chloramphenicol for plasmid transformation and outgrowth. For protein expression, cells were diluted in Terrific broth (Each 1 L medium contains 12g tryptone, 24g yeast extract and 4 ml glycerol diluted in water), supplemented with K-Phos salt solution (170 mM KH2PO4 and 0.72mM K2HPO4) and Kanamycin and grown at 37°C until OD=0.4-0.5, 0.4 mM Isopropyl β-d-1-thiogalactopyranoside (IPTG) were added to induce protein expression at 16°C for 24 hours.

### Plasmids

For overexpression and gene deletion CTDNEP1, NEP1R1, MAN1 and PPM1A, cDNAs and sgRNAs were cloned in a dual promoter lentiviral vector as described in ^67^. Constructs of CTDNEP1 WT, CTDNEP1D67E, D69T mutant, NEP1R1 and PPM1A cDNAs were cloned into a lentiviral vector in which the protein expression was driven by the EF-1α promoter. Expression of MAN1 WT, MAN1 mutants and SMAD2 were doxycycline inducible and driven by cytomegalovirus (CMV) promoter. For recombinant CTDNEP1 protein expression used in the in vitro dephosphorylation assay, vector containing H14-SUMO was used, with a lac operon to control protein expression by IPTG.

### Lentivirus production

For gene transductions using lentiviruses, virus was produced using Lenti-X 293T cells in 24-well plates using TransIT LT1 (Mirus Bio LLC #MIR 2305) and second-generation packaging vectors pMD2.G and psPAX2 according to standard lentiviral production protocols.

### Generation of CRISPR/Cas9-mediated knockout cells

Cell lines were transfected with single bicistronic sgRNA-Cas9 plasmid using Mirus LT-1 according to the manufacturer’s protocol. On the next day, cells were selected using Puromycin (2 µg/mL; Gibco #A1113803) for 72 hours. To generate KO clones, cells were single cell sorted using a BD AriaFusion flow cytometer. The knockout status of the clones was confirmed via immunoblotting and genomic sequencing. For genomic sequencing of KO clones, genomic DNAs (gDNA) were extracted from cells using QIAamp® DNA Blood Mini Kit (QIAGEN #51104) according to manufacturer’s protocol. PCR primers targeting ∼250bp upstream and downstream of the sgRNA cut site were used to amplify the purified gDNAs, resulting PCR products were analyzed by Sanger sequencing.

### Co-immunoprecipitation

Cells were lysed in 1% Decyl Maltose Neopentyl Glycol (DMNG) (Anatrace #NG322) lysis buffer (50 mM Tris-HCl pH 7.5, 150 mM NaCl) containing cOmplete EDTA-free protease inhibitor cocktail (Roche #5056489001) and phosphatase inhibitor cocktail (PhosSTOP, Roche #04906837001). Lysates were rotated at 4°C for 120 min. Cell debris and nuclei were precipitated at 20,000 g at 4°C for 20 min. Postnuclear supernatants were incubated for 2 h with anti-HA magnetic beads (PierceTM, Thermo Fisher Scientific #88837), or anti-FLAG magnetic beads (Sigma-Aldrich #M8823) or anti-V5 magnetic beads (MBL #M167-11). After three times 10min washes in 0.1% DMNG washing buffer (50 mM Tris-HCl pH 7.5, 150 mM NaCl), proteins were eluted in 1× sample buffer for 15 min at 65°C. The eluate was transferred to a new Eppendorf tube and subsequently reduced using dithiothreitol (DTT) (Merck Life Science UK Limited #D9779).

### Mass spectrometry-based analysis of immunoprecipitates

Triplicates of CTDNEP1 WT-3 × FLAG and 3 × V5-MAN1 WT and their corresponding empty vector control cell lines were subjected to cell lysis and co-immunoprecipitation as described in the previous session. When eluted, samples were resuspended in 25 µL 1 x SDS sample buffer (5% SDS, 50 mM Triethylammonium bicarbonate (TAEB), pH 7.55). Disulfide bonds were reduced using 20 mM TCEP for 15 mins at 47°C. After cooling down to room temperature, samples were alkylated using 20 mM CAA in the dark for 15 minutes and 10% volume of 12% phosphoric acid was added to acidify the samples. S-trap binding buffer (90% methanol in 100 mM TAEB, pH 7.5) was added to acidified, denatured samples to a final volume of 190 µL and the resulting solution was loaded onto S-Trap micro spin columns (ProtiFi), with a maximum of 150 µL of sample per load. Loaded spin columns were centrifuged at 4,000x g for 1 minute and this step was repeated until the entire sample was loaded onto a spin column. S-Trap columns were washed 5 × with S-trap binding buffer (90% methanol in 100 mM TAEB, pH 7.5) and the columns were transferred to 2 mL low protein binding Eppendorf tubes. For Trypsin/Lys-C digestion, 25 µL of digestion solution (50 mM TAEB, pH 8.0), containing 2 µg of Trypsin/Lys-C mix (Promega V5071) was added to each S-trap column, and the columns were incubated for 3 hours at 47°C on a ThermoMixer. Peptides were eluted with 30 µL of 50 mM TAEB, followed by 30 µL of 0.2% formic acid and 40 µL of 50% acetonitrile in 0.2% formic acid. Peptides were dried for 4 hours at 37°C in a vacuum centrifuge and samples were stored at −80°C until further analysis.

### Mass spectrometry analysis

Peptides were dissolved in 2% acetonitrile containing 0.1% trifluoroacetic acid, and each sample was independently analysed on an Orbitrap Fusion Lumos Tribrid mass spectrometer (Thermo Fisher Scientific), connected to an UltiMate 3000 RSLCnano System (Thermo Fisher Scientific). Peptides (1 µg) were injected on a PepMap 100 C18 LC trap column (300⍰μm ID⍰×⍰5 mm, 5⍰μm, 100⍰Å) followed by separation on an EASY-Spray nanoLC C18 column (75⍰μm ID ×⍰50 cm, 2⍰μm, 100⍰Å) at a flow rate of 250⍰nL/min. Solvent A was water containing 0.1% formic acid, and solvent B was 80% acetonitrile containing 0.1% formic acid. The gradient used for analysis of proteome samples was as follows: solvent B was maintained at 2% for 5⍰min, followed by an increase from 2 to 35% B in 120⍰min, 35-90% B in 0.5⍰min, maintained at 90% B for 4⍰min, followed by a decrease to 3% in 0.5⍰min and equilibration at 2% for 10⍰min. The Orbitrap Fusion Lumos was operated in positive-ion data-dependent mode. The precursor ion scan (full scan) was performed in the Orbitrap in the range of 400-1,600⍰m/z with a resolution of 120,000 at 200⍰m/z, an automatic gain control (AGC) target of 4⍰×⍰10^5^ and an ion injection time of 50⍰ms. MS/MS spectra were acquired in the linear ion trap (IT) using Rapid scan mode after high-energy collisional dissociation (HCD) fragmentation. An HCD collision energy of 30% was used, the AGC target was set to 1⍰×⍰10^4^ and dynamic injection time mode was allowed. The number of MS/MS events between full scans was determined on-the-fly to maintain a 3⍰s fixed duty cycle. Dynamic exclusion of ions within a⍰±⍰10⍰ppm m/z window was implemented using a 35⍰s exclusion duration. An electrospray voltage of 2.0⍰kV and capillary temperature of 275°C, with no sheath and auxiliary gas flow, was used.

### Mass spectrometry data analysis

All spectra were analysed using MaxQuant 1.6.10.43 ^68^ and searched against a SwissProt homo sapiens fasta files (containing 42,371 database entries with isoforms, downloaded on 2021/02/24). Peak list generation was performed within MaxQuant and searches were performed using default parameters and the built-in Andromeda search engine. The enzyme specificity was set to consider fully tryptic peptides, and two missed cleavages were allowed.

Oxidation of methionine and N-terminal acetylation were allowed as variable modifications. Carbamidomethylation of cysteine was allowed as a fixed modification. A protein and peptide false discovery rate (FDR) of less than 1% was employed in MaxQuant.

iBAQ intensities were used for data analysis. Briefly, the data was filtered to remove proteins that matched to a contaminant or a reverse database, which were only identified by site, which were not quantified in every sample, or which contained less than 2 unique peptides. iBAQ intensity values were log2 transformed.

### Immunoblotting

For immunoblotting, samples were incubated with 1 x SDS sample buffer (1x sample buffer: 67 mM Tris-HCl (pH6.8), 2% SDS, 10% glycerol, 0.067% BromophenolBlue) with DTT at 65°C for 10 min, separated by SDS–PAGE (Bio-Rad) and proteins were transferred to PVDF membranes (Bio-Rad).

Membranes were blocked in 5% Milk or bovine serum albumin (BSA, Sigma, A9418) in phosphate-buffered saline (PBS, Thermo Fisher Scientific #D8537-500mL) containing 0.1% Tween20 (Sigma #P1379-500mL) (PBS-Tween20) buffer and then probed with primary antibodies overnight at 4°C on a shaker. After three washes with PBS buffer, secondary antibodies were performed at RT for 1 hr either in 5% Milk or BSA in PBS-Tween20 buffer and subjected to another three washes with PBS. Membranes were developed by ECL (Western Lightning ECL Pro, PerkinElmer), and visualized using an Amersham Imager 600 (GE Healthcare Life Sciences).

### Immunofluorescence assay

HeLa cells were seeded onto EprediaTM round coverslips (Thermo Scientific Menzel #17294914). Expression of wild-type and mutant MAN1 was induced with 1 µg/ml doxycycline for 24 hours. For experiments required TGFβ treatment, 2ng/mL TGFβ3 were treated for 1 hour before fixation. Cells were then fixed with 4% methanol-free Paraformaldehyde (PFA) for 10 min at RT, followed by three PBS washes. Fixed cells were permeabilized in 0.2% Triton-X100 in PBS for 10min, followed by 1 hr blocking in blocking buffer (3% BSA, 0.2% Triton-X100 in PBS). Primary antibodies cocktails were prepared in blocking buffer. For primary antibody staining, coverslip was placed onto pre-spotted antibodies cocktail (40 μl/coverslip) on clean parafilm and incubated for 1 hr underneath home-made aluminum foil-covered moisturized chamber. Coverslips were washed three times with blocking buffer and secondary antibody staining was performed as was done for primary staining. Coverslips were then incubated in DAPI-containing PBS for 3 min, washed with PBS, and subsequently mounted onto glass slides in non-hardening mounting media and sealed with nail polish.

### Time-course analysis of SMAD2 localization

To analyze SMAD2 localization, cells were incubated with 2 ng/mL TGF-β3 for 1 h, followed by PBS washing for three times before adding 10 µM SB431542 (cell signaling #14775) inhibitor, cells were fixed either before adding inhibitor or after 1 h, 3 h, 7 h and 19 h inhibitor treatment. Samples were processed for immunofluorescence as described above.

### Confocal fluorescence microscopy

Fixed cells on slides after immunofluorescence preparation were imaged using an inverted Zeiss 880 microscope fitted with an Airyscan detector using ZEN black software. The system was equipped with Plan-Apochromat ×63/1.4-NA oil lens, with an immersion oil (Immersol W 2010, Carl Zeiss; refractive index of 1.518). 488 nm argon and 405, 561, and 633 nm solid-state diode lasers were used to excite fluorophores. Z-sections of images were collected. The oil objective was covered with an immersion oil (ImmersolT 518F, Carl Zeiss) with a refractive index of 1.518. Microscopy images with CZI file format were analyzed using ImageJ (1.53c, bundled with Java 1.8.0_172) software. Image quantification was performed by CellProfiler (4.2.1).

### pSMAD2 and pSMAD1 dephosphorylation kinetics assay

For testing the kinetics of pSAMD2 and pSAMD1 dephosphorylation, 8 × 10^4^ cells were seeded for each time point in the previous day. The next day, cells were incubated with 2 ng/mL TGF-β3 (Cell signaling #8425 and #10858) or 20 ng/mL BMP-4 (Peprotech 120-05) for 1 h. After stimulation with different cytokines, cells were then subjected to PBS washing for three times before adding 10 µM SB431542 (cell signaling #14775) or 0.1 µM LDN 193189 (Cambridge Bioscience HY-12071A-10mg) and incubated for different time.

For SB431542 inhibitor, cells were collected after 30min, 1 h and 2 h treatment and lysed with 1 × sample buffer. For LDN 193189 treatment, cells were lysed after 20 min, 40 min and 60 min incubation. Cell lysates were separated with gel electrophoresis followed by immunoblotting analysis.

### Inhibition of TGF-β with 1D11 neutralising antibody

Inhibition of TGF-β ligands was performed with the neutralising antibody, 1D11 (BioX-Cell # #BE0083), and isotype-matched IgG1 monoclonal control antibody (BioX-Cell #BE0057) were used at 30, 150 and 300 μg/mL respectively to treat HeLa parental cells or CTDENP1 KO cells with indicated time. Cell lysates were separated with gel electrophoresis followed by immunoblotting analysis.

### RT-qPCR

Total RNA was extracted from HeLa Parental, PPM1A KO, MAN1 KO, CTDNEP1 KO and NEP1R1 KO cell lines using Monarch® Total RNA Miniprep Kit (NEB T2010S). RT-qPCR was performed using the Luna Universal qPCR Master Mix (NEB M3003L) on a QuantStudio three Real-Time PCR System (Applied Biosystems) according to the manufacturer’s instructions. Primers used for amplifying the specific regions of p21, p15 and PAI1 are shown in (Table S2). The individual-specific gene transcript levels in the KO cell lines are normalised to GAPDH transcriptional level and presented as fold change relative to HeLa parental control, determined using the comparative CT (ΔΔCT) method. All reactions were carried out in two to three biological replicates with each replicate analyzed in three technical replicates.

### Cell proliferation assay and Flow cytometry

For EdU incorporation assay to assess cell proliferation, cells were seeded in 6-well plates for 24h prior to 30 minutes incubation with 10µM EdU (Abcam, #ab146186). Cells were then harvested, washed with PBS, and fixed with 70% Ethanol overnight. Fixed cells were pelleted by centrifuging at 1000xg for 5min, washed with PBS, and permeabilized with PBSTri-BSA (PBS, 0.1% Triton X-100, 1% BSA) on ice for 15min. Permeabilized cells were pelleted and washed twice with PBST-BSA (PBS, 0.1% Tween20, 1% BSA). EdU present in the cells were stained by Click-IT reaction (2mM CuSO4, 10mM Sodium Ascorbate, 10µM Alexa Fluor™ 555 Azide, Triethylammonium Salt (ThermoFisher Scientific #A20012) in PBS) at room temperature for 1h in dark. After Click-IT reaction, cells were pelleted and washed twice with PBST-BSA. DAPI staining was then performed in DAPI containing solution (PBS, 0.1% BSA, 1mg/mL RNAse A (ThermoFisher Scientific, # EN0531), 1µg/mL DAPI (BD Bioscience, #564907)) for 1h in room temperature in dark. Next, cells were analyzed using a BD LSRFortessa X-20 flow cytometer. For each condition, 30,000 cells were measured and FACS data was analyzed using FlowJo v10.

### Recombinant CTDNEP1 protein purification

Recombinant CTDNEP1 (46-244) WT and CTDNEP1D67E, D69T mutant containing an N-terminal His14-Sumo tag and a C-terminal myc tag were expressed in BL21-CodonPlus(DE3)-RIPL Competent cells (Agilent Technologies #230280). Bacterial cells were grown in terrific broth (supplemented with K-Phos salt solution and Kanamycin) at 37°C until OD600=0.4∼0.5. Protein expression was then induced by 0.4 M Isopropyl β-d-1-thiogalactopyranoside (IPTG) at 16°C for 24 hours. Cells were harvested by centrifugation at 4000 rpm for 10min and washed once with wash buffer (50 mM Tris-HCl (pH 8.0), 500 mM NaCl, 30 mM imidazole, 1 mM PMSF, 1.8 mM PepA). Cells were lysed by 1 mg/ml lysozyme + 0.05 mg/ml DNaseI in WB, incubating at room temperature for 30 min. After incubation, cell lysates were sonicated on ice by a probe sonicator (Soniprep 150), with amplitude set at 10-15 microns and on/off pulse of 30 seconds duration for 5 times. The lysate was cleared by centrifugation at 4000 rpm for 10 min. Cleared lysates were subjected to ultracentrifugation in Ti45 tubes at 40,000 rpm for 45 min at 4°C to separate a crude membrane fraction. Supernatants were collected and incubated with Ni-NTA Agarose beads (Thermo Scientific, HisPurTM #88222) overnight at 4 °C. After incubation, the material was transferred to a 20 ml gravity column and beads were washed by gravity flow with 20 column volumes of WB. Proteins that were bound on the Ni-NTA beads were eluted by elution buffer (50 mM Tris-HCl (pH 8.0), 500 mM NaCl, 300 mM imidazole).

Eluted protein was pooled and loaded onto a Superdex 200 Increase 10/300 GL column (GE #28-9909-44) equilibrated with buffer (20 mM HEPES (pH 7.4), 200 mM NaCl). Peak fractions were collected and snap frozen until use. To obtain His-SUMO protein as the control of the in vitro phosphatase, an aliquot of purified His-SUMO-CTDNEP1 WT was immobilized on Ni-NTA beads in 10 mL of WB for 1 hour at 4 °C. 1 µM of Ulp1 was added to the beads and incubated further for 2 hours at 4 °C. The material was transferred to a 20 mL gravity column and beads were washed by gravity flow with 10 column volumes of WB. His-SUMO protein bound on the Ni-NTA beads was eluted by elution buffer and snap frozen.

### In vitro phosphatase assay

Lentiviral plasmid encoding SMAD2 with N-terminal 3 × HA tag was transduced into a HeLa MAN1 and CTDNEP1 double KO clone. Cells were treated with 1µg/ml doxycycline for 24 hours to induce HA-SMAD2 expression, followed by 20 ng/mL TGFβ3 treatment for 1 hour to obtain pool of phospho-SMAD2. Cells were lysed with 1% DMNG followed by anti-HA immunoprecipitation as described in previous method section. In vitro dephosphorylation assay was applied to HA-pSMAD2 beads with recombinantly expressed and purified 100 ng and 1 µg CTDNEP WT and wild mutant protein at 37 °C for 10 min. For quenching the assay, the beads were incubated with 1 × SDS sample buffer and boiled at 65°C for 20 min.

### Western blot quantification

Western blot data was quantified using Image Studio software (Li-Cor Ver5.2) and graphs were plotted using Prism (GraphPad). Representative images of at least three independent experiments are shown. Student t test was used for statistical analysis and error bars represent the standard deviation.

### Quantification of SMAD2 localization

Nucleo:Perinuclear ratio of SMAD2 intensity was quantified by CellProfiler (4.2.1). Raw microscopic images of SMAD2 and DAPI were imported to ImageJ (1.53c, bundled with Java 1.8.0_172) software to acquire maximum projection of the z-stack images. Maximum projection image files of SMAD2 and DAPI were imported to CellProfiler. DAPI staining was used as the indicator of the nuclear area, and SMAD2 staining was used to indicate the whole cell area.

The cytoplasmic area was determined by subtracting the nuclear area from the whole cell area. Within the cytoplasmic area, perinuclear area was defined by a 10-pixel ring surrounding the nucleus. Integrated intensity of SMAD2 in the nuclear area and perinuclear area was measured and the ratio was calculated in logarithms for statistical analysis. One-way ANOVA test and graph plotting were performed using GraphPad Prism 10. For each sample, more than 100 cells were analyzed, which were obtained from at least three independent experiments and at least three different fields per sample from each experiment. Error bars represent the standard error of the mean of three replicates.

### Quantification of EdU staining

Flow cytometry data was analyzed by FlowJo v10. For each sample, the EdU-positive cells were gated from a total of 30,000 cells, and the percentage of EdU+ cells from two independent replicates were compared between control cells and the KO cells in GraphPad Prism 10 using one-way ANOVA. Error bars represent the standard deviation.

### Data and Code Availability

The mass spectrometry proteomics data have been deposited to the ProteomeXchange Consortium via the PRIDE ^69^ partner repository with the dataset identifier PXD051056. The published article includes all processed data generated or analyzed during this study as Supplemental Information.

All original code is available in this paper’s supplemental information. Any additional information required to reanalyze the data reported in this paper is available from the lead contact upon request.

## Notes

### Competing Interest Statement

The authors have declared no competing interest.

