## Supplementary figures and images for "Suppression of TGF-β/SMAD signaling by an inner nuclear membrane phosphatase complex"

### Fig S1

#### Supplemental Figure 1

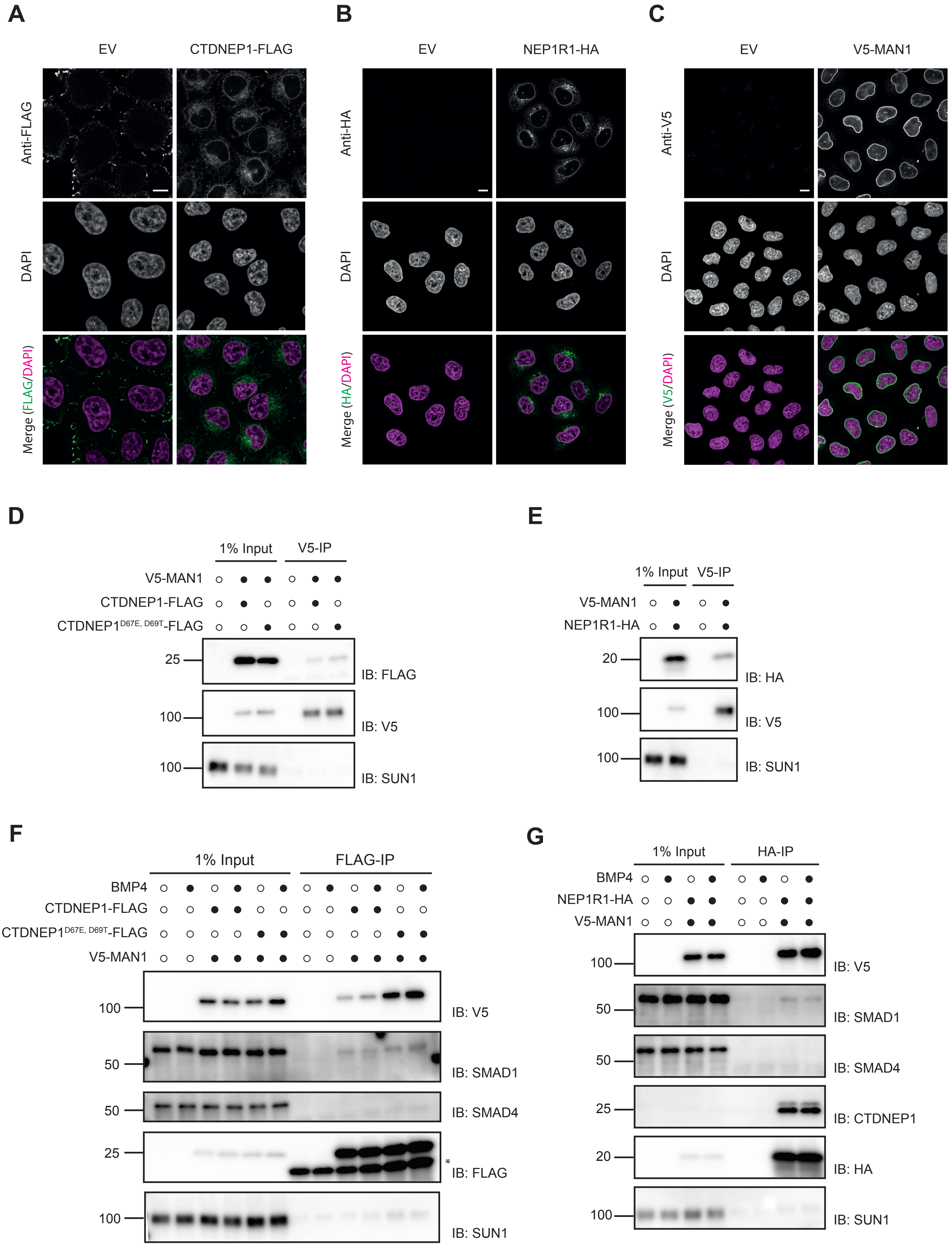

### Fig S2

#### Supplemental Figure 2

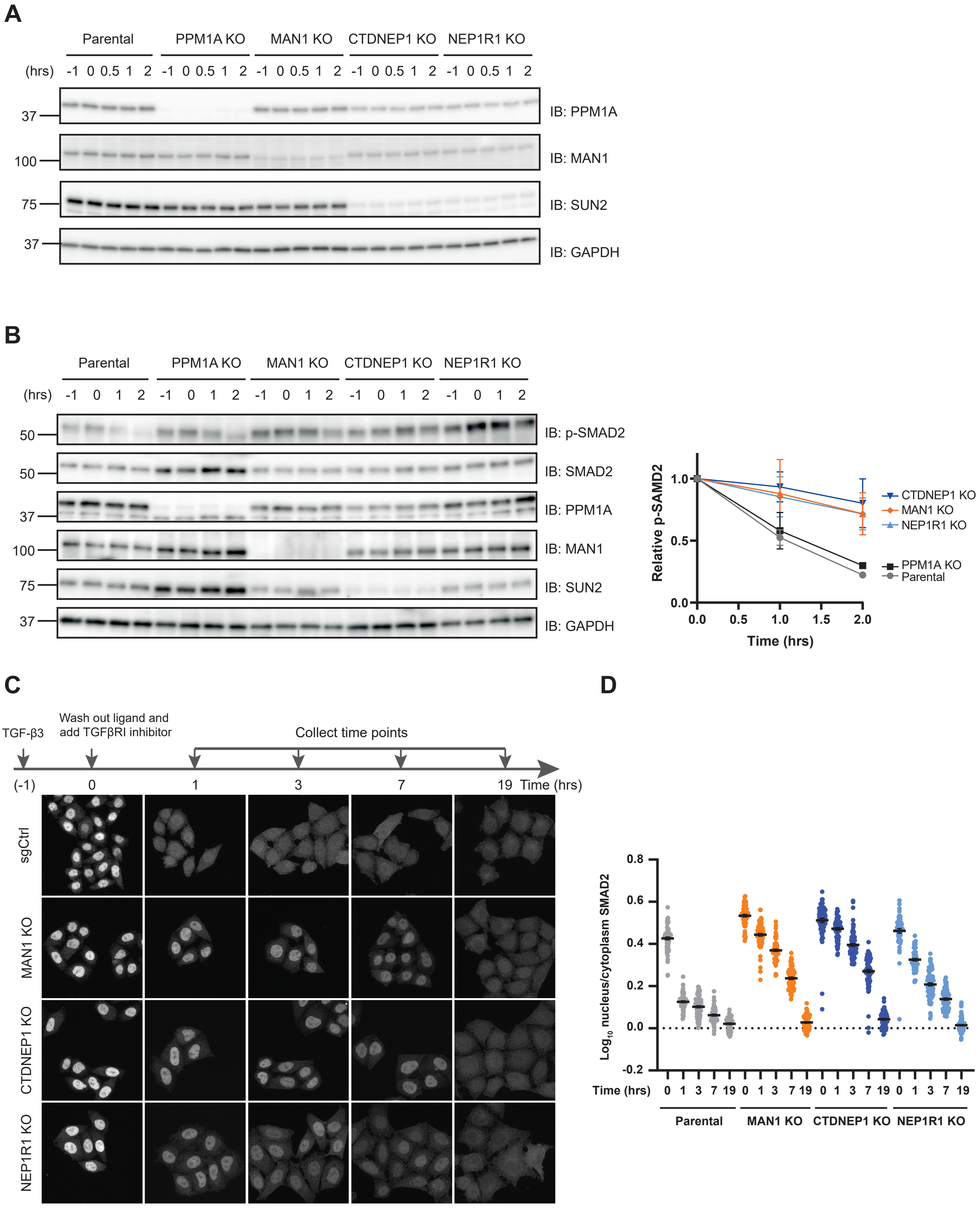

### Fig S3

#### Supplemental Figure 3

**A**

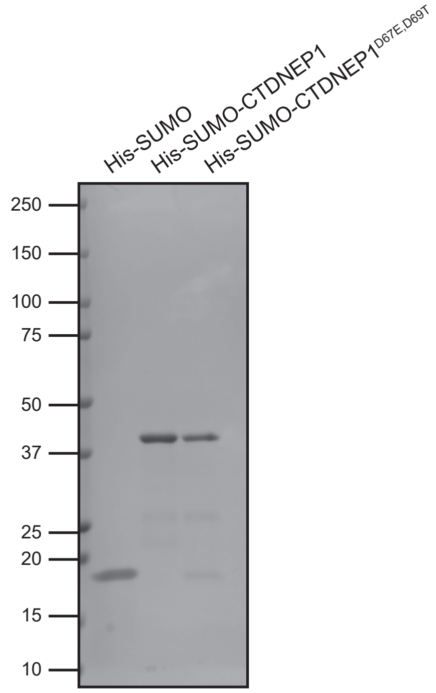


**B**

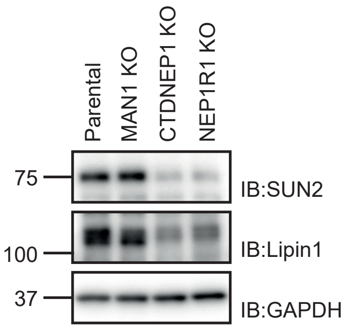

### Fig S4

#### Supplemental Figure 4

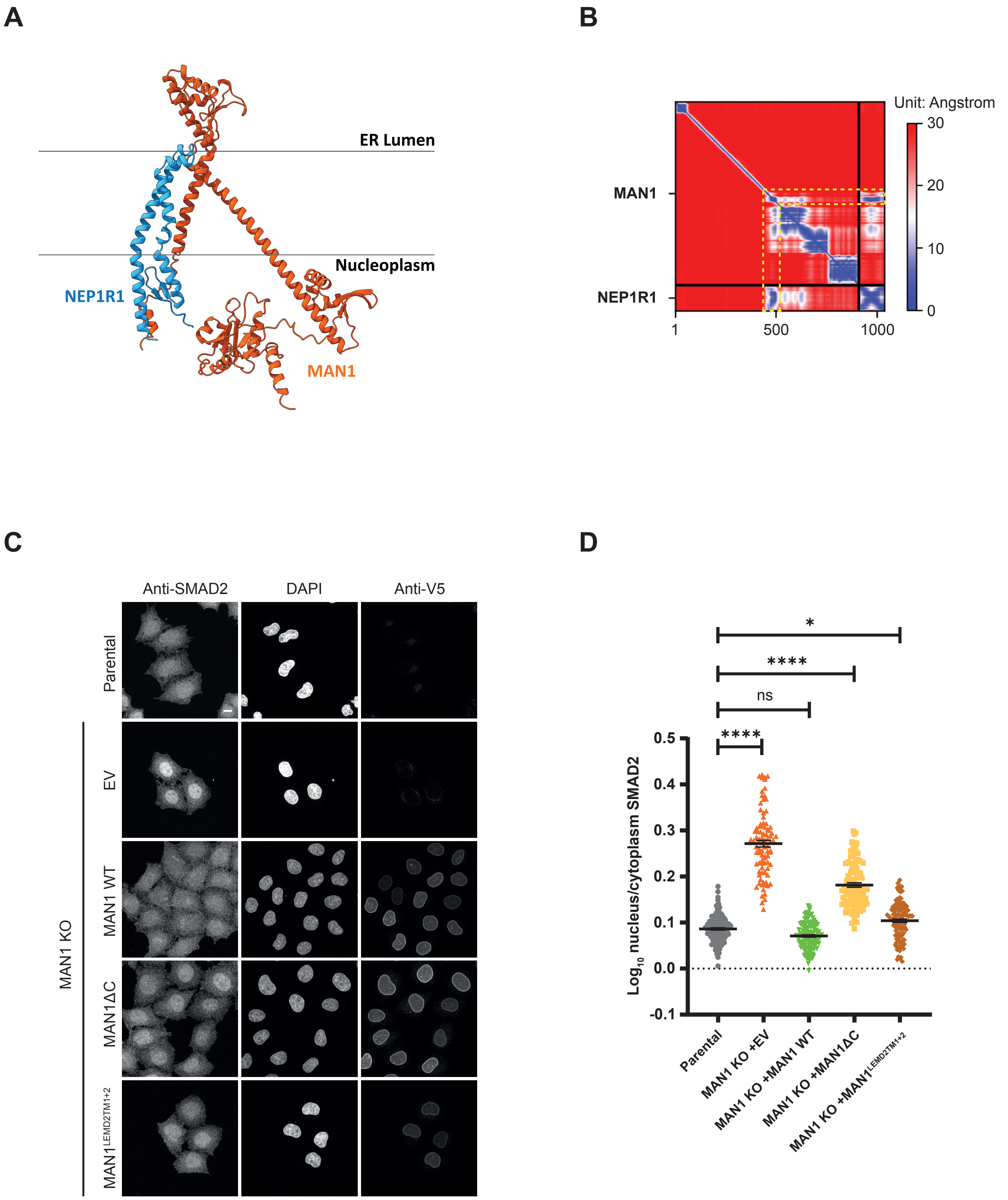

### Fig S5

#### Supplemental Figure 5

A

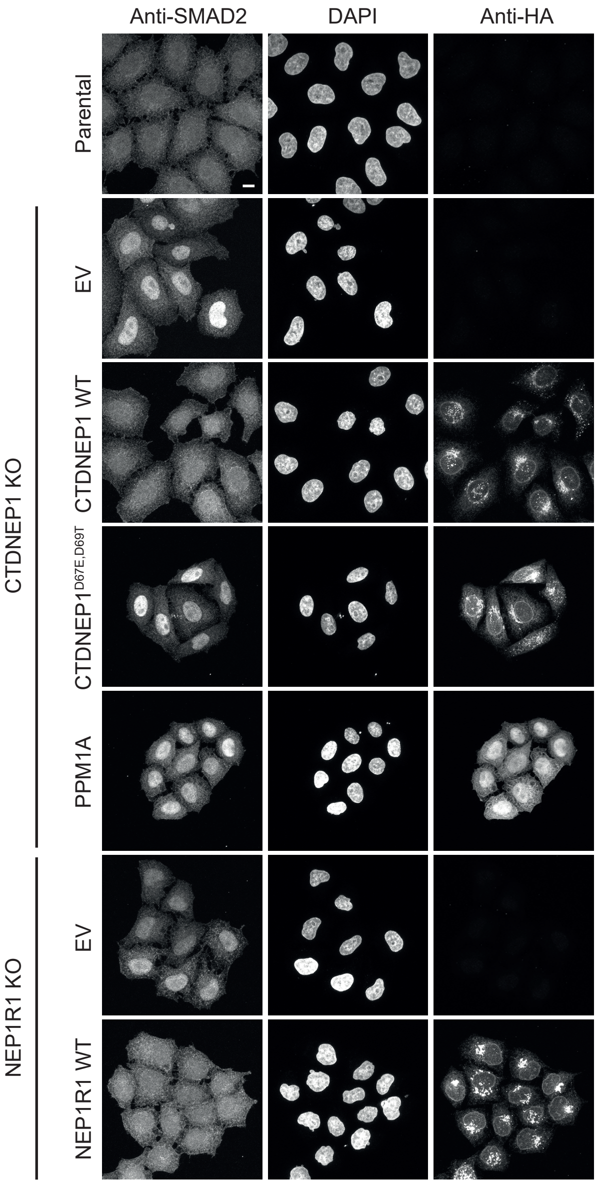


B

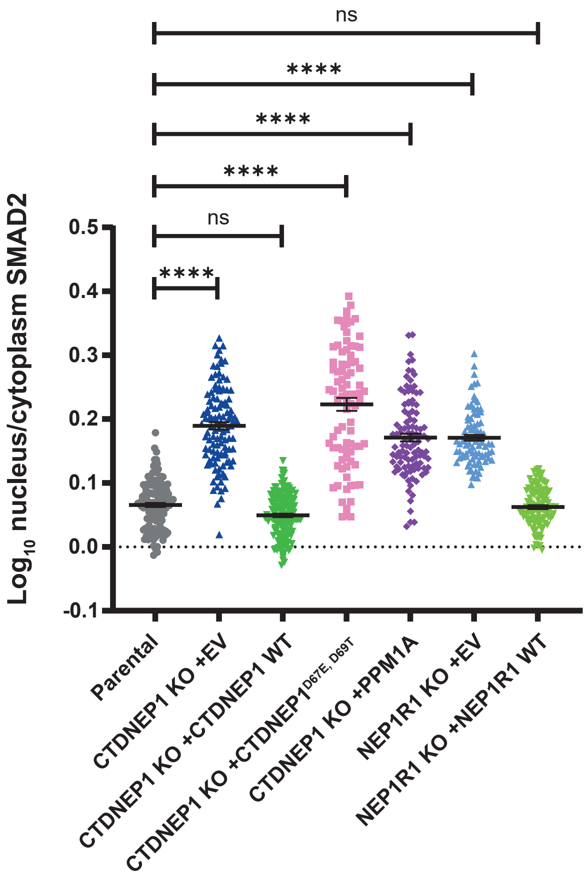


C

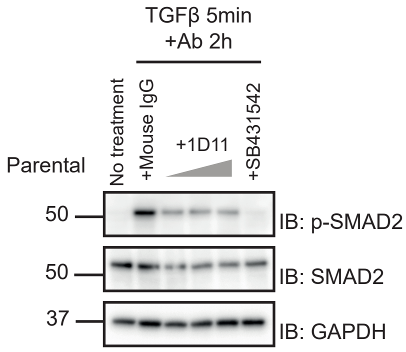


D

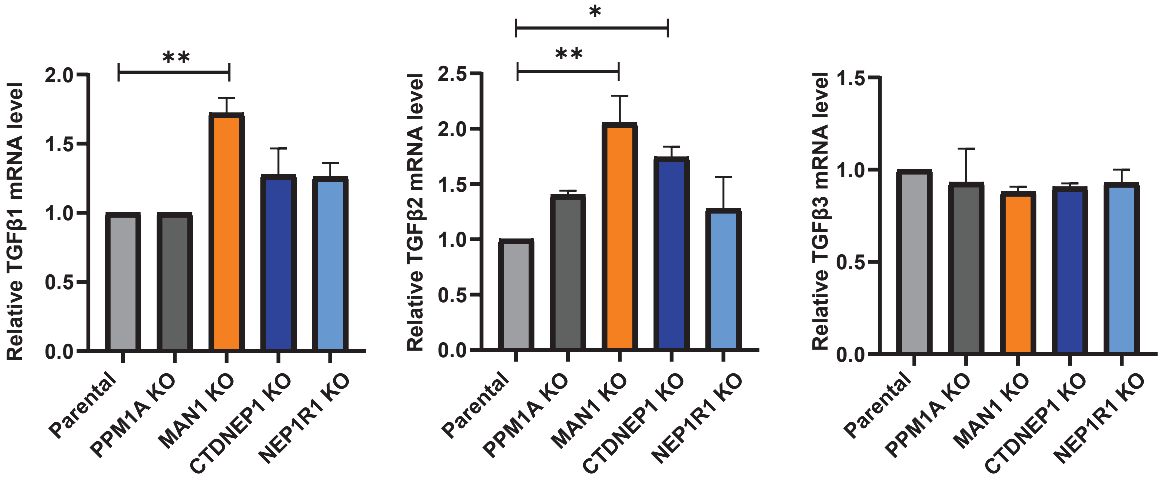
